# ER stress and lipid imbalance drive embryonic cardiomyopathy in a human heart organoid model of pregestational diabetes

**DOI:** 10.1101/2023.06.07.544081

**Authors:** Aleksandra Kostina, Yonatan R. Lewis-Israeli, Mishref Abdelhamid, Mitchell A. Gabalski, Brett D. Volmert, Haley Lankerd, Amanda R. Huang, Aaron H. Wasserman, Todd Lydic, Christina Chan, Isoken Olomu, Aitor Aguirre

## Abstract

Congenital heart defects constitute the most common birth defect in humans, affecting approximately 1% of all live births. The incidence of congenital heart defects is exacerbated by maternal conditions, such as diabetes during the first trimester. Our ability to mechanistically understand these disorders is severely limited by the lack of human models and the inaccessibility to human tissue at relevant stages. Here, we used an advanced human heart organoid model that recapitulates complex aspects of heart development during the first trimester to model the effects of pregestational diabetes in the human embryonic heart. We observed that heart organoids in diabetic conditions develop pathophysiological hallmarks like those previously reported in mouse and human studies, including ROS-mediated stress and cardiomyocyte hypertrophy, among others. Single cell RNA-seq revealed cardiac cell type specific-dysfunction affecting epicardial and cardiomyocyte populations, and suggested alterations in endoplasmic reticulum function and very long chain fatty acid lipid metabolism. Confocal imaging and LC-MS lipidomics confirmed our observations and showed that dyslipidemia was mediated by fatty acid desaturase 2 (FADS2) mRNA decay dependent on IRE1-RIDD signaling. We also found that the effects of pregestational diabetes could be reversed to a significant extent using drug interventions targeting either IRE1 or restoring healthy lipid levels within organoids, opening the door to new preventative and therapeutic strategies in humans.

## INTRODUCTION

Congenital heart disease (CHD) constitutes the most common type of congenital defect in humans^1^. Pregestational diabetes (PGD, defined as diabetes of the mother before and during the first trimester of pregnancy), regardless of whether it is T1D or T2D, is one of the most outstanding non-genetic factors contributing to CHD^2^ and is present in a significant, growing population of diabetic female patients of reproductive age^2^. Newborns from mothers with PGD have increased risk of CHD (up to 12%), versus ∼1% in the normal population (12-fold increase)^2–5^. PGD is hard to manage clinically due to extreme sensitivity of the developing embryo to glucose oscillations and constitutes a critical health problem for the mother and the fetus. The prevalence of PGD-induced CHD is increasing significantly due to the ongoing diabetes epidemic, and particularly affects underserved populations^6–8^. Preventative and therapeutic interventions are urgently needed to tackle this health problem.

In the last decades, significant efforts have been devoted to understanding the molecular pathology of PGD-induced CHD using animal models^9, 10^, leading to the identification of increased ROS production, abnormal lipid metabolism, epigenetic reprogramming, and mitochondrial stress among others^4, 11–16^. The molecular links and the overall contribution of these abnormalities to the different phenotypes observed in PGD-induced CHD remain elusive, and despite these efforts, no advances have been translated to the clinical setting. A core issue is that it remains unclear to what extent rodent models recapitulate abnormalities present in human PGD-induced CHD, given critical species differences in heart size, cardiac physiology and electrophysiology, and bioenergetics^17–19^. For example, there are major species differences between mice and humans in how the heart utilizes glucose, lactate, ketone bodies and fatty acids. Furthermore, previous rodent models (and most in vitro cell models) commonly rely on overtly aggressive diabetic conditions (∼25-50 mM blood glucose in diabetic mice vs ∼11-33 mM in human diabetic patients)^9^, which might lead to exaggerated cytotoxic phenotypes not relevant to the human clinical condition under study. Clinical practice largely precludes studies of PGD-induced CHD in humans, or severely limits them, thus direct studies of human embryos are not viable options. This is understandable given the care that pregnant mothers need, particularly if they suffer from potential increased risk to their fetus. However, the result is significantly limited access to human tissues for research of early-stage disease and mechanisms of PGD-induced CHD, forcing an overreliance on animal models and stalling progress to understand the condition.

Novel stem cell-based technologies have enabled the creation of engineered, highly complex, human organ-like 3D tissues *in vitro*, with properties that recapitulate the physiological setting to a significant extent^20–24^. These organoids are particularly useful to study unapproachable disease states in humans (e.g., early disease progression when symptoms are not yet present), or states for which animal models are not well-suited^25–28^. Organoids can also be powerful tools to verify in human models what we have learned in animal studies, complementing them. While organoids have been used to model a wide range of human tissues and conditions, their application to cardiovascular studies has been sorely lacking until very recently^25–27^. We recently established an advanced heart organoid model that recapitulates human heart development during the first trimester reliably or accurately^29, 30^, including critical steps such as chamber formation, vascularization, cardiac tissue organization and relevant cardiac cell types^22, 29, 31^.

In this report, we describe the application of our heart organoid model to the study of PGD-induced CHD. We found that heart organoids could be employed to model critical aspects of CHD and recapitulated hallmarks of PGD-induced CHD found in mice and humans. Interestingly, for the first time, we found a new cause of CHD seemingly specific to the human setting. Maternal diabetes causes significant stress in the ER, leading to disrupted very long chain fatty acid (VLCFA) lipid metabolism due to degradation of FADS2 mRNA via the IRE1- RIDD pathway. VLCFAs are critical signaling and structural components of cardiac cells during development so it is not surprising that their deficiency can lead to CHD. Finally, we found that restoring VLCFA levels by either inhibiting ER stress or exogenous dietary administration of VLFCAs can greatly reduce the deleterious effects of PGD, opening the door to new clinical interventions to prevent and treat these disorders.

## RESULTS

### Human heart organoids faithfully recapitulate pathological hallmarks of pregestational diabetes-induced congenital heart disease

To model pregestational diabetes conditions in the embryonic heart, we developed high glucose and high insulin medium with concentrations of these molecules similar to those previously reported in patients^32, 33^. hHOs (human heart organoids) were exposed to normoglycemic medium (NHOs) or hyperglycemic medium (PGDHOs) from their assembly at day 0 of differentiation and maintained in these conditions throughout *in vitro* development until assayed for a series of biochemical and cellular hallmarks of PGD-induced CHD described before^12, 13, 15, 16^. To determine the presence of cardiomyocyte hypertrophy, a well-described effect of embryonic diabetic cardiomyopathy^34–38^, we dissociated day 14 NHOs and PGDHOs into single cell suspensions and performed immunofluorescence staining in individual cardiomyocytes for TNNT2. A significant enlargement of cardiomyocytes was observed in PGDHOs compared to NHOs (**Fig. 1A, B**), resulting in a 27±9.5% increase in cardiomyocyte size (**Supp. Fig. 1A**). Mitochondrial staining using the molecular probe Mitotracker revealed abnormal mitochondrial morphology and mitochondrial swelling in PGDHOs compared to NHOs (**Fig. 1C, D; Supp. Fig. 1B, C**), suggesting mitochondrial disfunction due to diabetic conditions and pointing potential metabolic phenotypes. It has been has been previously described in murine PGD models^10, 33^ that pregestational diabetes induces oxidative stress in the developing embryo^16, 39^. Analysis for production of reactive oxygen species (ROS) in NHOs and PGDHOs revealed an increase in cellular ROS in PGDHOs compared with controls (**Fig. 1E, F**). We also characterized the dynamic expression of key developmental transcription factors *(HAND1, HAND2, NKX2-5, TBX5* and *GATA4*) (**Fig. 1G-J; Supp. Fig. 1D**) in NHOs and PGDHOs between days 0 and 14. NHOs displayed a clear transition in the expression of the first heart field (FHF) marker *HAND1* which was highly expressed between days 2 and 8, and the expression of the second heart field (SHF) marker *HAND2* which was highly expressed from days 8 onwards. Both heart field markers were downregulated in PGDHOs compared to NHOs at critical time points. *NKX2-5* and *TBX5* both showed overexpression in the PGDHOs which was not present in their control counterparts, the first showing an early overexpression on day 6 and the latter showing an overexpression on days 12 and 14. *GATA4* also presented abnormal expression over time (**Supp. Fig. 1D**). These data were consistent with previous observations of developmental transcription factor dysregulation in PGD conditions^27^. We also investigated the expression of glucose transporters *SLC2A1* and *SLC2A4* (**Supp. Fig. 1E, F**). NHOs demonstrate a clear transition between the glucose transporter *SLC2A1* (highly expressed in the fetal heart) and the glucose transporter *SLC2A4* (highly expressed in the adult heart) between days 6 and 8. On the other hand, PGDHOs show downregulation of *SLC2A1* in the early days of differentiation, hinting at possible dysfunction in glycolysis and glucose transport (**Supp. Fig. 1E**). Normal hHOs typically exhibit a well-developed primitive vascular network around the myocardial tissue during differentiation^29^. NHOs and PGDHOs showed drastic differences in the formation of this vascular network. The endothelial cell marker PECAM1 revealed a sophisticated plexus of vascular endothelial tissue covering large portions of the control NHOs (**Fig. 1K**). Vascularization in PGDHOs appeared less organized than in NHOs, showing fewer regions of PECAM1 staining and lack of an interconnected network (**Fig. 1K**). To compare functional properties of control and diabetic hHOs, calcium transient analysis by live organoid imaging was conducted. The beating frequency of the control organoids was ∼120 beats per minute (bpm) compared to ∼60 bpm in diabetic organoids (**Fig. 1L, M**). Fetal heart rate can range from 110 to over 160 bpm^34^, with slower fetal heart rate associated with poor pregnancy outcomes^35^. PGDHOs also showed a higher frequency of arrhythmic events when compared to NHOs (**Fig. 1L**). Overall, these data demonstrate substantial morphological and molecular changes specific to PGDHOs that are consistent with observations of fetal tissue in animal and human models^12, 13, 15, 16, 37^.

**Figure 1.**
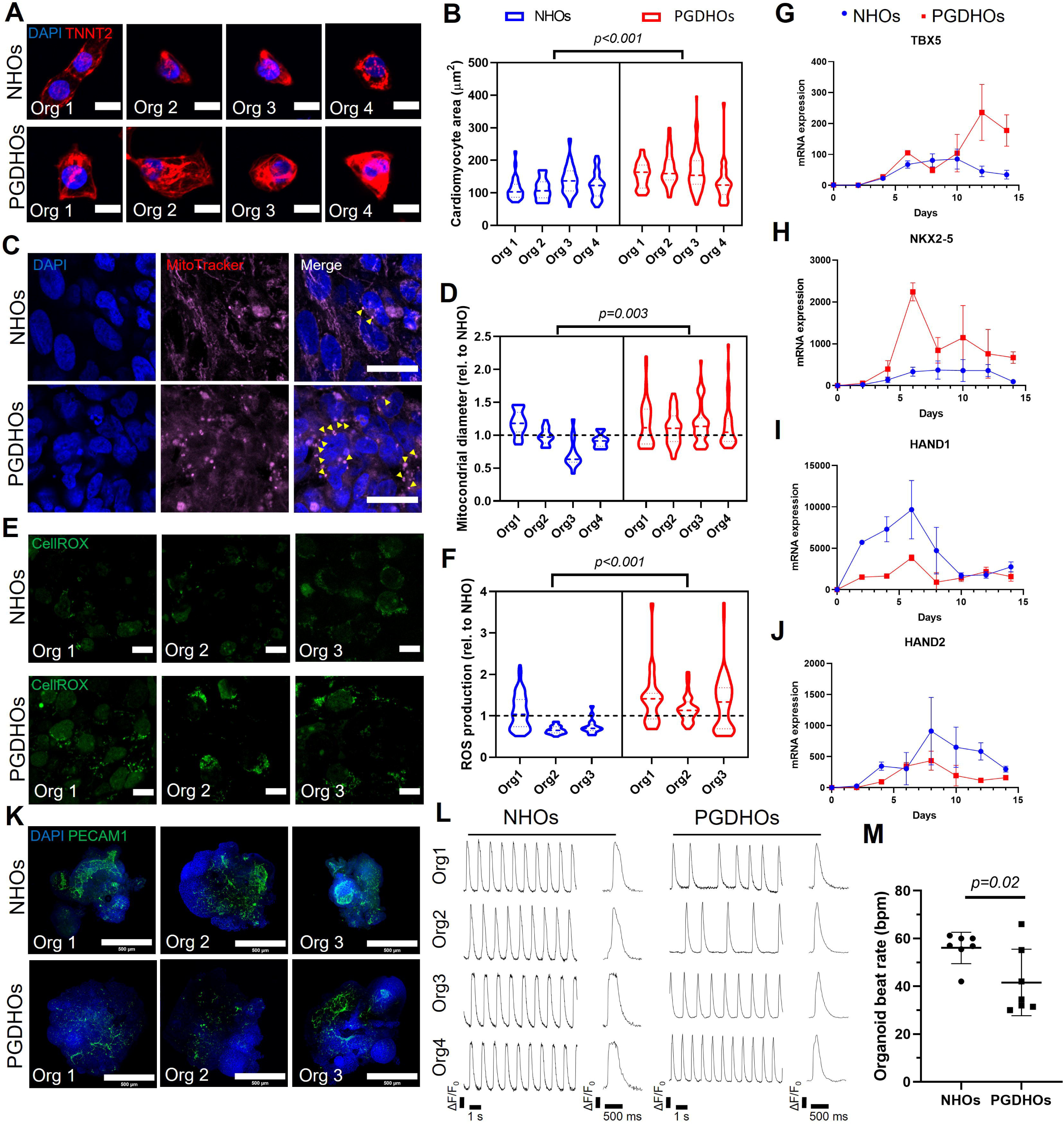
Human heart organoids recapitulate phenotypic hallmarks of pregestational diabetes in the developing human heart. **A**, Immunofluorescence images of cardiomyocytes from NHOs and PGDHOs stained with the cardiomyocyte marker TNNT2 (red) and the nuclear marker DAPI (blue); scale bar: 10 µm. **B**, Quantification of cardiomyocyte area from NHOs and PGDHOs; n=4 organoids, n=145 cardiomyocytes for NHOs and n=193 cardiomyocytes for PGDHOs, nested t-test. **C**, Immunofluorescence of mitochondria using MitoTracker showing mitochondrial swelling in PGDHOs; arrowheads indicated swollen mitochondria, scale bar: 10 µm. **D**, Quantification of mitochondrial swelling in NHOs and PGDHOs; n=4; nested t-test. **E**, Live ROS imaging employing CellROX Green; scale bar: 10 µm. **F**, Quantification of ROS content in NHOs and PGDHOs; n=3; nested t-test. **G–J**, Time course qRT-PCR gene expression analysis of key developmental transcription factors from day 0 to day 14 of differentiation; n=9 (3 biological replicates of 3 pooled organoids). **L**, Calcium transient live imaging using Fluo-4 from NHOs (left) and PGDHOs (right), and **M**, Quantification of beats per minute in NHOs and PGDHOs; n=7; value = mean ± SD, unpaired t-tests.

### Single cell RNA sequencing reveals cardiac cell type-specific responses to PGD in heart organoids

To explore the effects of PGD on cardiac lineage specification within heart organoids, we performed single-cell RNA sequencing on NHOS and PGDHOS at day 15 of differentiation. Computational analysis identified 6 main clusters representing distinct cardiac cell populations in both NHOs and PGDHOs. These clusters included cardiomyocytes, immature cardiomyocytes, epicardial cells, endothelial cells, cardiac fibroblasts and a mixed population of mesenchymal cardiac progenitors which were still poorly committed (labelled as others) (**Fig. 2A, B**). The main effects of PGD were a significant decrease in the number of cardiomyocytes (30% in NHOs vs 23% in PGDHOs) and a very large increase in epicardial cells (5% in NHOs vs 24% in PGDHOs) (**Fig. 2C**), hinting at dysregulation of cardiomyocyte and epicardial differentiation^36^. The effects of PGD on classical cardiomyocyte and epicardial markers can be observed in t-SNE plots (**Fig. 2D, E**). A summary of other key markers related to other cell populations and their changes in PGD can be found in **Suppl. Fig A-C**. Interestingly, a sinoatrial node-related subcluster was also observed within NHOs but not in the PGDHO group (**Supp. Fig. 2C**). A summary of the top 20 most significant differentially regulated genes in epicardial cells identified dysregulation of epicardial transcription factor expression (*TBX18*), Wnt and Notch signaling (*WNT2B*, *SFRP2*, *DLK1*) and other developmental regulators, such as retinoic acid signaling (*ALDH1A2*) in PGD conditions (**Fig. 2F**). In the case of cardiomyocytes, analysis of the top 20 differentially expressed genes identified sarcomeric and cytoskeletal gene expression alterations in PGD, particularly in genes frequently associated with familial cardiomyopathies and hypertrophy (e.g., *CSRP3, SMPX, MYOZ2*). Genes involved in fatty acid metabolism and ER function were also significantly dysregulated in cardiomyocytes (*FITM1, HSPB3*). Interestingly, when all clusters were considered together, the top hit was also a strongly downregulated gene involved in ER protein synthesis (*EMC10*). To confirm previous observations pointing to alterations in FHF and SHF formation, we produced combined t-SNE plots for key FHF markers (*HAND1*, *HCN4*, *TBX5*) and key SHF markers (*HAND2*, *ISL1*, *TBX1*) (**Supp. Fig. 2D**). Interestingly, cell populations from NHOs that showed expression of FHF markers cells were predominantly in the CM and immature CM clusters while many of the SHF marker-expressing cells appeared throughout the clusters in myocyte and non-myocyte cell populations. This observation is in agreement with the precardiac organoids derived from mouse ESCs demonstrating the development of non-myocyte cells from SHF progenitor cells^37^.

**Figure 2.**
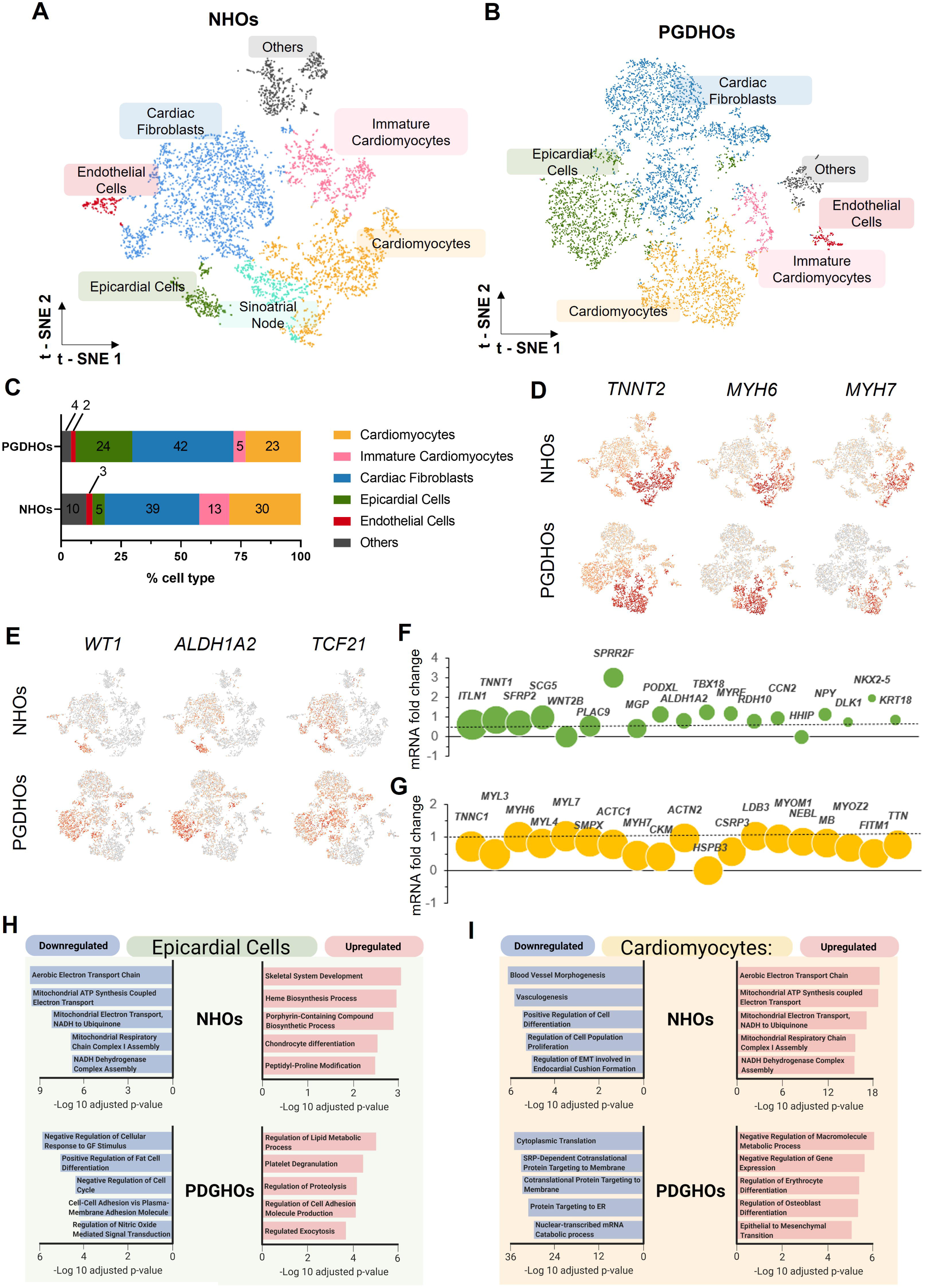
scRNA-seq reveals cardiac cell type-specific responses to PGD in human heart organoids. **A**, t-SNE plot showing 5,951 cells from 4 pooled dissociated NHO organoids, **B**, t-SNE plot of 7,463 cells from 4 pooled dissociated PGDHO organoids, distributed by k-means clustering. Each cluster was named based on the expression of key genes in the respective cell populations. **C,** Cardiac cell composition in NHOs and PGDHOs expressed as percentage of cells. **D**, t-SNE plots showing key markers for cardiomyocytes and (**E**) and epicardial cells. **F**, Bubble plot with top DEGs specific for epicardial cells and (**G**) cardiomyocytes. Size of bubble corresponds to p-value between NHOs and PGDHOs. **H**, Gene ontology analysis of biological processes associated with significantly expressed genes in the epicardial cluster and (**I**) the epicardial cluster (**H**).

To better understand the molecular differences between heart organoids grown in healthy conditions and PGD conditions, we performed gene ontology for differentially expressed genes (DEGs) for the main identified clusters (**Fig. 2A, B**). DEG analysis revealed 373, 289, and 315 significantly expressed DEG for the cardiac fibroblasts, epicardial and endothelial cell clusters, respectively (**Supp. Fig. 2E**). Gene ontology (GO) for downregulated DEGs in epicardial cells of NHOs revealed markers of mitochondrial function while upregulated DEGs in PGDHOs were associated with regulation of lipid metabolic processes (**Fig. 2H)**. Bioinformatic analysis revealed 379 DEGs in the CM cluster, 223 and 156 significantly expressed genes distinct to NHOs and PGDHOs, respectively **(Supp. Fig. 2E)**, with a focus on mitochondrial respiration and ATP synthesis in NHOs, while downregulated DEGs were associated with vessel morphogenesis, cell differentiation and proliferation (**Fig. 2I**). In contrast, PGDHOs revealed upregulation of genes related to negative regulation of metabolic processes and differentiation of non-myocytes, and downregulation of translation processes and protein targeting to the endoplasmic reticulum (ER) (**Fig. 2I**). These data again pointed at developmental defects in cardiomyocytes under PGD conditions related to ER function and lipid metabolism. Gene ontology (GO) analysis for the most significantly expressed genes from the cardiac fibroblasts and endothelial cells clusters is presented in **Supp. Fig. 2F, G**, showing the similarities between clusters from NHOs and PGDHOs. Overall, our scRNA-seq analysis revealed critical differences in transcriptomic profiles of normoglycemic and diabetic organoids affecting early heart development, particularly cardiomyocyte and epicardial populations, and pointed to potential dysregulation of ER and lipid metabolisms as important contributors in the development of PGD-induced CHD.

### PGD triggers ER stress and VLCFA dyslipidemia in human heart organoids

Previous work has shown a strong association between ER stress and cardiomyopathies^40^. Our data suggested that ER stress might also be a factor in PGD-induced CHD. We decided to explore the potential connection between ROS production and ER stress in cardiomyocytes as a result of diabetic conditions^16, 39^. hHOs were co-stained with CellROX Green and ER Tracker Red, a highly selective dye for the ER, and imaged under normoglycemic and PGD conditions. PGDHOs exhibited increased cellular ROS predominantly accumulated to the ER (**Fig. 3A, B**), suggesting potential ER dysfunction. Interestingly, ER stress has not been described in animal models of PGD and may be a human-specific phenotype of interest for therapeutic and preventative interventions. Inositol-requiring enzyme 1 (IRE1, gene name *ERN1*) is the main evolutionarily conserved sensor for ER stress. IRE1 phosphorylation within the ER membrane induces the unfolded protein response (UPR) through at least two pathways, one of them involves the splicing of *XBP1* mRNA and activation of downstream transcription effectors. The second pathway is named regulated IRE1-dependent decay (RIDD), and targets mRNAs for degradation in an XBP1-independent manner^41^. Confocal imaging for activation of IRE1 demonstrated an increase in the ratio of the phosphorylated isoform (IRE1p) when compared to its unphosphorylated counterpart, confirming PGDHOs have higher ER stress than NHOs (**Fig. 3C, D, Supp. Fig. 3A**). Another molecular phenotype identified in our scRNA-seq analysis was lipid metabolism. LC-MS lipidomic profiling revealed significant differences between PGDHOs and NHOs affecting very long chain fatty acid (VLCFAs) trafficking and synthesis (**Fig. 3E**). Organoids were grown in fully defined medium, so no exogenous sources of VLCFAs or other lipids are available aside from essential fatty acids present in the medium. All other fatty acids present are therefore synthesized by the organoids to be used intracellularly or to be secreted into the extracellular space. The intracellular levels of two important omega-3 fatty acids, EPA and DPA, were significantly higher in PGDHOs (**Fig. 3E),** while the levels of DHA (another important omega-3 fatty acid synthesized from EPA and DPA), were significantly lower in the medium for PGDHOs (**Fig. 3E)**. This data showed that the PGD conditions resulted in lipid dysregulation in VLCFAs, particularly in omega-3 fatty acids, which have been previously shown to be instrumental during heart development^40–42^. The mechanistic implication of these changes in cardiac physiology remains unclear, since omega-3 fatty acids can be used as building blocks in cell membranes^42^, signaling lipids (docosanoids)^43^ and antioxidants^44^.

**Figure 3.**
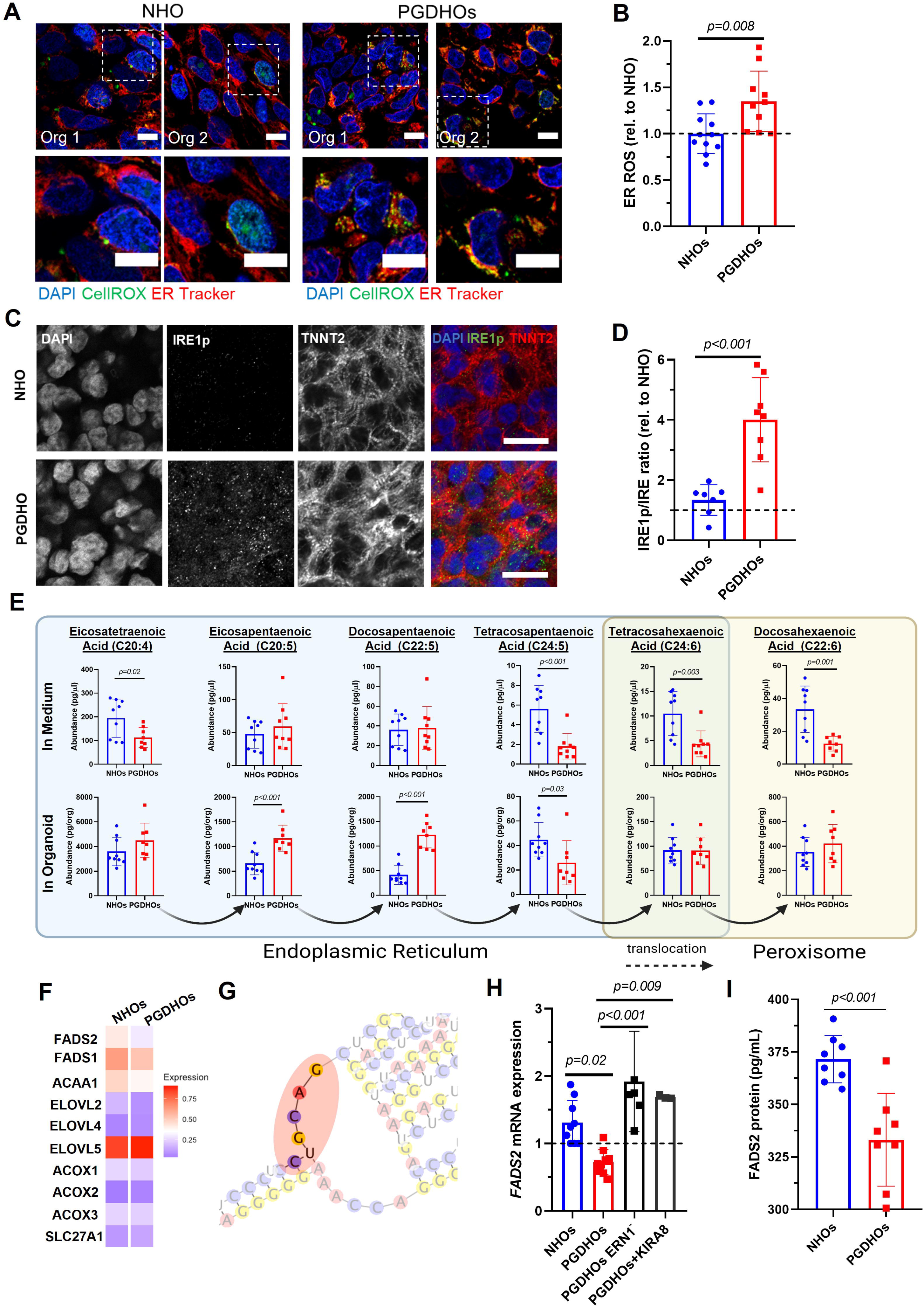
Pregestational diabetes conditions trigger ER stress and VLCFA lipid imbalance in human heart organoids. **A**, Immunofluorescence images of day 14 NHOs and PGDHOs showing colocalization of ROS (CellROX, green) and ER marker (ER Tracker, red); nuclear marker DAPI (blue); scale bar=10 µm. **B**, Quantification ROS localized in the ER; n=10 organoids per condition; value = mean ± SD, unpaired t-tests. **C**, Immunofluorescence images of day 14 NHOs and PGDHOs stained for phosphorylated IRE1 (IRE1p, green), cardiomyocyte marker TNNT2 (red) and nuclear marker DAPI; n=6; scale bar=20 µm. **D**, Quantification of the ratio of phosphorylated IRE1 to unphosphorylated IRE1 compared to NHOs, measured by immunofluorescence image analysis; n=7 for NHOs, n=8 for PGDHOs; value = mean ± SD, unpaired t-tests. **E,** LC-MS lipidomic analysis for VLCFA concentrations from day 15 organoids and their corresponding medium; n=9 organoids per condition, value = mean ± SD, unpaired t- test, *p<0.05. **F**, Heatmap representing expression level of key enzymes involved in LCFA and VLCFA biosynthesis. **G**, Predicted secondary *FADS2* mRNA structure. **H**, qRT-PCR gene expression analysis of *FADS2*; n=6 and n=3 biological replicates of 3 pooled organoids for ERN1 knockdown and KIRA8 respectively; value = mean ± SD. **I**, FADS2 protein measurement by ELISA; n=8 organoids per condition; value = mean ± SD.

Next, we attempted to find a link between ER stress and disrupted VLCFA metabolism. The ER plays a significant role in the synthesis of lipid precursors, such as long chain fatty acid (LCFAs) necessary for the synthesis of more complex lipids, and we hypothesized ER stress might interfere with this balance. We started by looking at IRE1-induced activation of the UPR, which can lead to apoptosis or re-establishment of protein homeostasis. To determine if IRE1 activation was causing apoptosis we used a transgenic iPSC line expressing Flip-GFP, a non-fluorescence engineered GFP variant that turns fluorescent upon caspase activation and apoptosis^45^. Flip-GFP organoids cultured either under control or diabetic conditions exhibited no fluorescence changes indicating that there is no significant PGD-induced apoptosis (**Supp. Fig. 3B**). Doxorubicin-treated NHOs and PGDHOs were used as a positive control for apoptosis (**Supp. Fig. 3B)**. Furthermore, analysis of UPR-specific genes, either induced by IRE1 or some other ER stress-sensor (ATF6, PERK) demonstrated absence of UPR pathway activation (**Supp. Fig. 3C).** IRE1 signaling through XBP1 splicing also remained unchanged between NHOs and PGDHOs (**Supp. Fig. 3D**). We then decided to investigate whether the IRE1-RIDD pathway was activated due to ER stress, and if that was the process affecting lipid biosynthesis.

RIDD has been reported to preferentially induce degradation of ER-localized mRNA^46^ and most steps of endogenous fatty acid biosynthesis take place in the smooth ER. We performed gene expression analysis for 10 ER-localized lipid biosynthesis enzymes in NHOs and PGDHOs and found increased expression of *ELOVL5* (ELOVL Fatty Acid Elongase 5) along with decreased expression of *ELOVL2* (ELOVL Fatty Acid Elongase 2), *ACAA1* (3-Ketoacyl-CoA thiolase) and desaturases *FADS1* (Fatty Acid Desaturase 1) and *FADS2* (Fatty Acid Desaturase 2, also known as D6D) in PGD conditions (**Fig. 3F)**. Delta-6 desaturase (D6D) encoded by the *FADS2* gene catalyzes the key initial rate-limiting step of VLCFA biosynthesis before exporting FAs to the peroxisome (tetracosapentaenoic acid to tetracosahexaenoic acid) and is a main determinant of VLCFA levels. Alterations in D6D activity alter fatty acid profiles and are associated with metabolic and inflammatory diseases^47–50^. LC-MS analysis had revealed significant differences in this metabolic step under the control of D6D both in organoids and medium (**Fig. 3E).** Interestingly, the FADS2 mRNA possesses four RIDD consensus sequences for degradation in its 3’ and 5’ UTR regions (5’-CUGCAG-3’ motif located in the loop portion of a hairpin structure) (http://www.geneious.com/) (**Supp. Fig. 3H)**, and at least one of them is in a clear hairpin loop **(Fig 3G)**. To confirm that *FADS2* mRNA downregulation is IRE1-RIDD-dependent we analyzed *FADS2* expression in a knockdown *ERN1* (IRE1) iPSC line and in the presence of IRE1 mono-selective inhibitor KIRA8. *ERN1* knockdown PGDHOs exhibited restored levels of *FADS2* expression comparable with NHOs and confirming the dependence of *FADS2* expression on IRE1-RIDD activation (**Fig. 3H)**. The same effect was found when IRE1 activity was inhibited using the specific inhibitor KIRA8 (**Fig. 3H**) in the absence XBP1 splicing (**Supp. Fig. 3G).** D6D protein abundance was measured by ELISA and was found to be significantly lower in PGDHOs compared to NHOs, confirming the pathologic phenotype **(Fig. 3I)**. Expression of *FADS1*, *ACAA1* and *ELOVL2* in heart organoids with *ERN1* knockdown was comparable with levels in PGDHOs, suggesting these enzymes are not regulated by IRE1-RIDD and confirming the importance of FADS2 in this setting (**Supp. Fig. 3E-G)**. Taken together, these results show that PGD causes ROS-induced ER stress and IRE-RIDD activation leading to delta-6 desaturase deficiency in diabetic heart organoids. This in turn affects VLCFA biosynthesis and causes a crucial VLCFA imbalance during early cardiac development.

### Reducing ER stress mitigates deleterious effects of PGD in human heart organoids

To ameliorate the phenotypic effects of PGD in the developing heart, and the resulting ROS overproduction and ER stress, we tested several therapeutic compounds based on our mechanistic findings involving IRE1 and omega-3 fatty acids. These included the chemical chaperone tauroursodeoxycholic acid (TUDCA), known to reduce oxidative and ER stress ^43–45^, a combination of omega-3 fatty acids (DHA, DPA, EPA) designed to provide antioxidant resilience and relieve lipid imbalance and the antidiabetic compound sapropterin (BH4)^49, 50^, which has been previously described as a potential mitigating agent for PGD-induced CHD in murine models^4, 50^. Treatment of PGDHOs with TUDCA, BH4 and omega-3 fatty acids all caused a significant reduction of phosphorylated IRE1 in PGDHO organoids (**Fig. 4A).** The ratio of IRE1p to IRE1 was significantly lower in all treated organoids as well, suggesting a reduction in IRE1 signaling activation (**Fig. 4B**). Co-staining of organoids with CellROX Green and ER Tracker revealed reduced stress due to ROS both in the ER and cytosol for BH4 and omega-3 fatty acids treatments, but not for TUDCA (**Fig. 4C, D**). Immunofluorescence imaging of individual cardimyocytes from dissociated organoids revealed that the treatment of PGDHOs with TUDCA, BH4 and omega-3 fatty acids resulted in a significant reduction in cardiomyocyte hypertrophy, suggesting an amelioration in the hypertrophic response (**Fig. 4E, F**). Notably, treating PGDHOs with TUDCA, BH4, or omega-3 fatty acids affected mRNA levels of *FADS2* (**Fig. 4G**), likely due to the reduction in IRE1 signaling. **Fig. 4H** shows a general mechanistic schematic summarizing our findings in the context of PGD-induced CHD. Overall, the tested therapeutic agents were able to decrease all the tested hallmarks of PGD-induced CHD, confirming our mechanistic hypothesis and validating the findings in mouse models and BH4. This is a potentially important finding for clinical translation, as it suggests that dietary supplementation (e.g., omega-3 fatty acids) of diabetic mothers in the first trimester of pregnancy might be able to prevent or treat early stages of PGD-induced CHD with no known drawbacks.

**Figure 4.**
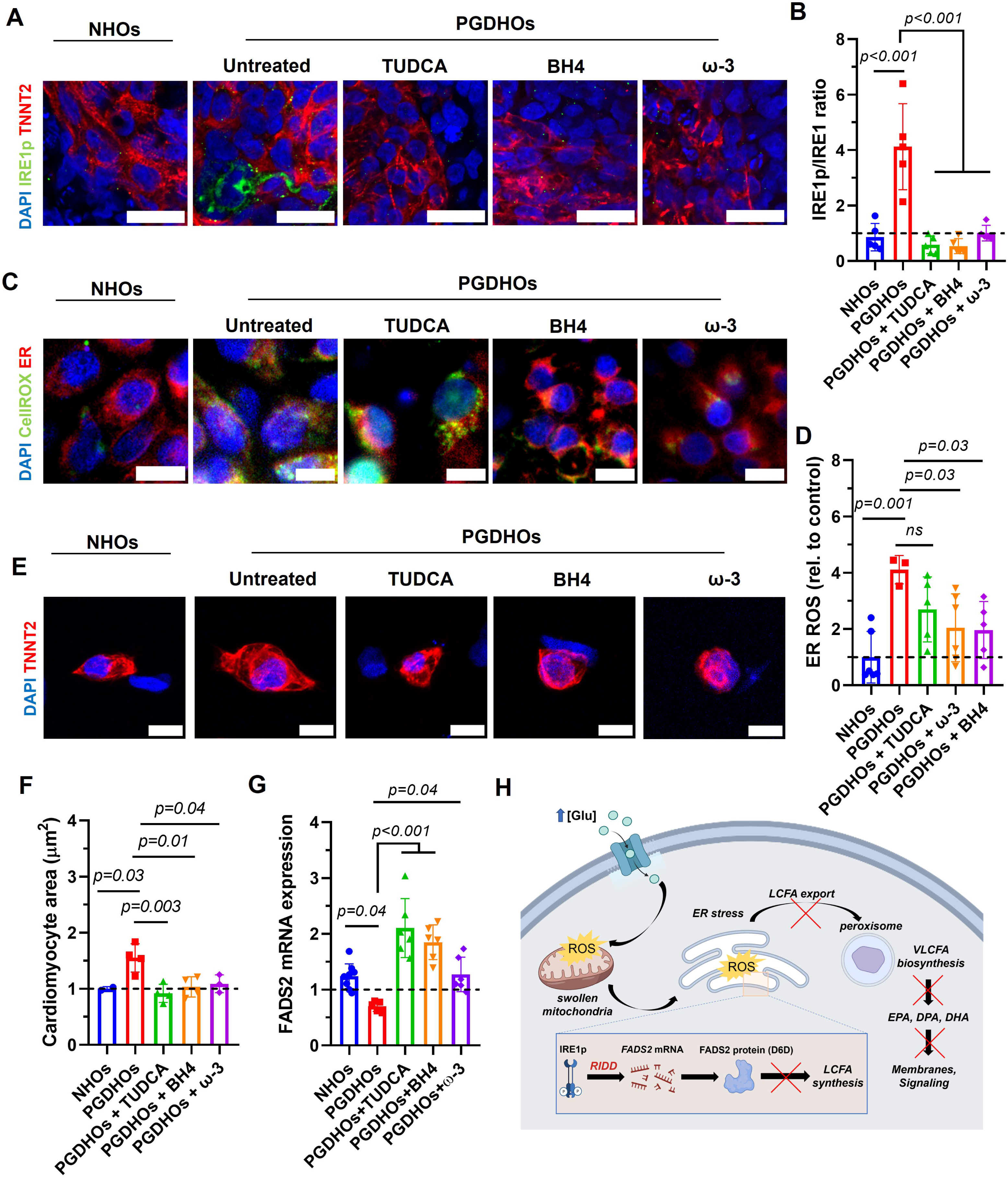
Strategies to reduce ER stress and lipid imbalance mitigate the deleterious effects of pregestational diabetes in human heart organoids. A, Immunofluorescence images of day 14 NHOs and PGDHOs treated with TUDCA, BH4, or omega-3 fatty acids, stained with phosphorylated IRE1 (IRE1p, green), cardiomyocyte marker TNNT2 (red) and nuclear marker DAPI (blue); n=6; scale bar: 25 µm. **B**, Quantification of the ratio phosphorylated IRE1 to unphosphorylated IRE1, measured by immunofluorescence image analysis; n=6; value = mean ± SD, one-way ANOVA. **C**, Immunofluorescence images of ROS (CellROX, green) and ER marked (ER Tracker, red) in day 14 NHOs and PGDHOs; n=7; scale bar = 10 µm. **D**, Quantification of ROS content compared to NHOs; n=6; value = mean ± SD, one-way ANOVA. **E**, Immunofluorescence images of cardiomyocytes dissociated from day 14 NHOs, PGDHOs; n=4; scale bar = 10 µm. **F**, Quantification of cardiomyocyte area compared to NHOs; n=4; value = mean ± SD, one-way ANOVA. **G**, qRT-PCR gene expression analysis of *FADS2* in NHOs and PGDHOs; n=6 biological replicates of 3 pooled organoids; value = mean ± SD. **H**, Schematic diagram summarizing the mechanism of PGD-induced CHD in human heart organoids.

## DISCUSSION

We utilized a novel and powerful heart organoid technology based on recent work^29, 51^ to the study of critical mechanisms of PGD-induced CHD in humans, and report, for the first time, that ER stress and VLCFA imbalance are critical factors contributing to these congenital disorders. Maternal diabetes is one of the most common causes of newborn CHD (up to 12% of newborns from diabetic mothers have some form of CHD^52^), yet the ability to study the etiology of these disorders in humans is greatly limited due to inaccessibility to human fetal hearts at crucial stages of development. The effects of PGD on CHD have been studied extensively in animal models, identifying many contributing factors including the dysregulation of lipid metabolism^53, 54^, an increase in oxidative stress^15, 55, 56^, and dysregulation of key transcription factors related to heart development^4, 11, 13, 14, 56–58^. However, the extent of the ability of these animal models to properly recapitulate human PGD-CHD abnormalities is limited by cross-species differences. hHOs provide a necessary platform on which to investigate the underlying mechanisms between the association of PGD and CHD in humans without the drawbacks of animal models.

To model the effects of PGD, hHO culture conditions were modified to accurately reflect reported physiological levels of glucose and insulin in normal and diabetic mothers for females with type I and type II pregestational diabetes. The larger size of diabetic hHOs suggested signs of cardiac hypertrophy, a first hallmark of maternal PGD^59^ on the embryo’s heart, which was confirmed by studying cardiomyocyte size in dissociated organoid cultures (**Fig. 1**). Furthermore, differences were noted with regards to ROS production and mitochondria of PGDHOs, revealing increased oxidative stress and mitochondrial swelling, also hallmarks of diabetic embryonic cardiomyopathies^36^. Mouse models of PGD have shown dysregulation of transcription factors key to heart development, including GATA4, GATA5, TBX5, and NKX2- 5^4, 11, 13, 14, 57, 58^. In line with this, key structural abnormalities were observed in PGDHOs, including disorganization of myocardial, epicardial and endothelial tissue, as well as disruption of atrioventricular specification and organization. Time course qPCR analysis revealed disparities in regulation of GATA4, NKX2-5 and TBX5 at different time points throughout development, highlighting the importance of fine transcription control during early development. Moreover, the specification of the FHF and subsequent SHF are crucial in the development and organization of the heart, and the observed downregulation of heart field markers in PGDHOs likely contributes to the structural malformation and under-representation of cardiomyocytes compared to NHOs. Similarly, the downregulation of the glucose transporter SLC2A1 during the early stages of differentiation suggests disruption in glycolysis may be crucial in the observed metabolic disfunction. These phenotypes were consistent with outcomes found in PGD-induced CHD in animal models and human clinical reports^15, 55, 56, 60^. PGDHOs also presented arrythmias and reduction in beat frequency as determined by calcium imaging, a phenotype that has not been described in diabetic embryonic cardiomyopathy but has been observed in neonatal rats from diabetic mothers^61^ and diabetic human adults, which typically exhibit reduced calcium pump activity leading to diastolic dysfunction^62^.

Single-cell transcriptomics provides a valuable tool to investigate cell-type and lineage specific changes between healthy and diabetic organoids. We performed scRNA-seq on day 15 PGDHOs and NHOs, revealing important differences in cardiac lineage specification. Most notably, a reduction in cardiomyocyte numbers, a significant expansion of epicardial tissue and the absence of a well-developed conductance system at early developmental stages (**Fig. 2**). Furthermore, a number of important candidate genes were identified for the first time in this context in both epicardial cells and cardiomyocytes, providing new hypothesis for future work and therapeutic interventions. In the case of the epicardium, a number of important growth factors, enzymes and developmentally relevant receptors were dysregulated (*WNT2B, SFRP2, ALDH1A2*) (**Fig. 2F**), suggesting the epicardium has a trophic effect on the rest of the heart at this stage^63^. *ITLN1*, a protein involved in glucose metabolism was also dysregulated^64, 65^. In the case of cardiomyocytes, major changes involved sarcomeric proteins (*TNNC1, MYL3, MYH6, MYL7* and others), but also other proteins involved in protein synthesis (*HSPB3*) and lipid metabolism (FITM1) (**Fig. 2G**). These changes were more radical than those observed in the epicardium overall (larger statistical significance, larger fold change), and prompted us to investigate the pathophysiology of PGD-induced CHD in cardiomyocytes in more detail.

Glycolysis constitutes a significant source of energy and intermediate metabolites in the embryonic heart, but fatty acid oxidation is also important for mitochondrial growth and cardiac maturation as the heart develops^28, 59, 60^. Our model confirmed heart organoids exposed to diabetic conditions exhibited increased accumulation of ROS (**Fig.1, 3**), but also showed that a significant portion of ROS was localized to the ER and could be impairing its function (**Fig. 3A**). Numerous studies have shown that elevated ROS produces ER protein oxidation^66^, leading to a condition known as ER stress. The ER plays also an important role in lipid metabolism as a significant part of lipid synthesis occurs in the smooth ER^67^. Out of the three sensor systems for ER stress that can activate the UPR (unfolded protein response), only IRE1 has been shown to be strongly associated with lipid metabolism (the other two being the ATF6 and PERK pathways)^68–72^. The lipid metabolic profiles of PGDHOs revealed a clear dysregulation of VLCFA synthesis particularly affecting omega-3 polyunsaturated fatty acids (PUFA). We and others have shown that VLCFAs are critical during heart development for structural and signaling reasons^73–75^, and the involvement of VLCFAs and ER stress has not been described before in PGD-CHD, either in mouse models or human in vitro systems. The endogenous synthesis of these PUFAs mostly takes place in the ER, and along with the observed ROS accumulation in the ER, suggested a major ER-induced dyslipidemia in PGDHOs. We could pinpoint this dysregulation to the targeted degradation of *FADS2*, a key lipid biosynthesis enzyme present in the ER, by the IRE1-dependent mRNA decay (RIDD) pathway, and thus for the first time we demonstrated a direct role of IRE1-RIDD in lipid metabolism dysregulation in PGD-induced CHD. Interestingly, IRE1-RIDD contribution has been described for several other cardiac pathologies^76, 77^, reinforcing our findings.

Our findings show that targeting the IRE1 pathway, or its downstream VLFCA dysregulation, could be significant for therapeutic targeting. VLCFA lipid imbalance is treatable by means of dietary supplementation (pregnant diabetic mothers, and their newborns, present better outcomes when supplemented with DHA)^78^. Alternatives to treat ER stress by chemically targeting IRE1 also exist and could be translated to the clinic if ER stress is a major driver of PGD-CHD. In an attempt to remedy the effects of ER stress, we tested several potentially therapeutic compounds on PGDHOs, including BH4, a recently identified molecule that can reduce the incidence of PGD-induced CHD in mice^11^, a mixture of omega-3 fatty acids (EPA, DPA, DHA), and TUDCA (an IRE1 modulator). Both BH4 and omega-3 PUFAs ameliorated diabetic phenotypes, and while TUDCA (an IRE1 modulator) did not significantly resolve the observed reactive oxygen species, it did ameliorate cardiomyocyte hypertrophy. All of the compounds also restored *FADS2* levels. These data suggest that while BH4 and Ω3 fatty acids help ameliorate the effects of PGD on the developing heart via reduction in reactive oxygen species, TUDCA’s therapeutic role may be more closely involved in transcription factor regulation. Lastly, the data suggests that the increased levels of phosphorylated IRE1 may be a key player in the mechanism behind PGD associated CHD.

In summary, we established the value of a heart organoid model of pregestational diabetes and CHD, and our findings unveiled a novel ROS-induced ER stress mechanism underlying critical VLCFA balance in PGD-CHD. These findings represent a new route for developing future preventative and therapeutic strategies, thus helping to reduce the incidence of CHD across the population and provide proof-of-concept of the utility of organoid models for human disease modeling and drug development.

## MATERIALS AND METHODS

### Stem cell culture

An in-house human iPSC line was used in this study (iPSC-L1). hiPSCs were validated for pluripotency, genomic instability, and potential contamination on a regular basis. For apoptosis experiments, a transgenic iPSC-L1 line expressing Flip-GFP^45^. hPSCs were cultured in Essential 8 Flex medium containing 1% penicillin/streptomycin (Gibco) on 6-well plates coated with growth factor-reduced Matrigel (Corning) in an incubator at 37°C, 5% CO_2_ until 60-80% confluency was reached, at which point cells were split into new wells using ReLeSR passaging reagent (Stem Cell Technologies). Accutase (Innovative Cell Technologies) was used to dissociate iPSCs for spheroid formation. After dissociation, cells were centrifuged at 300 g for 5 minutes and resuspended in Essential 8 Flex medium containing 2 µM ROCK inhibitor Thiazovivin (Millipore Sigma). hPSCs were then counted using a Moxi Cell Counter (Orflo Technologies) and seeded at 10,000 cells/well in round bottom ultra-low attachment 96- well plates (Costar) on day -2 at a volume of 100 µl per well. The plate was then centrifuged at 100 g for 3 minutes and placed in an incubator at 37°C, 5% CO_2_. After 24 hours (day -1), 50 µl of media was carefully removed from each well, and 200 µl of fresh Essential 8 Flex medium was added for a final volume of 250 µl/well. The plate was returned to the incubator for an additional 24 hours.

### Heart organoid differentiation and pregestational diabetes modeling

Pregestational diabetes was modeled as previously described^29, 51^. Briefly, diabetic conditions were simulated by using basal RPMI media with 11.1 mM glucose and 1.14 nM insulin and compared with control media containing 3.5 mM glucose and 170 pM insulin, henceforth referred to as diabetic and healthy culture media, respectively. The differentiation of the healthy or diabetic organoids was conducted as previously described^27^. Briefly, on day 0 of differentiation, 166 µl (∼2/3 of total well volume) of media was removed from each well and 166 µl of diabetic or healthy culture media without any insulin, containing CHIR99021 (Selleck) was added at a final concentration of 4 µM/well along with BMP4 at 0.36 pM (1.25ng/ml) and Activin A at 0.08 pM (1ng/ml), for 24 hours at 37°C, 5%. On day 1, 166 µl of media was removed and replaced with fresh culture media without insulin. On day 2, 166 µl of media was removed and replaced with fresh culture media minus insulin, containing Wnt-C59 (Selleck) for a final concentration of 2 µM Wnt-C59 in each well, and incubated for 48 hours at 37°C, 5%. On day 4, 166 µl of media was removed and replaced with fresh culture media without insulin. On day 6, 166 µl of media was removed and replaced with fresh culture media with respective diabetic and healthy insulin in the culture media. On day 7, a second 2 µM CHIR99021 exposure was conducted for 1 hour in fresh culture media at 37°C, 5%. Subsequently, media was changed every 48 hours until organoids were ready for analysis. For IRE1 inhibition organoids were treated with 1 µM mono-selective inhibitor KIRA8 (Selleckchem) for 5 days. Organoids were analyzed on day 14 unless otherwise indicated.

### shRNA-mediated lentiviral knockdown experiments

For lentivirus production, HEK293T cells were transfected with the shERN1-puro plasmid (VectorBuilder) and the packaging plasmids (pMD2; psPAX2) using lipofectamine with Plus reagent (Thermo). Lentivirus was added to iPSCs with 8□μg/ml polybrene (Fisher Scientific) and incubated overnight. Puromycin selection was carried out for ∼3–5 days. Surviving clones were collected, replated, and expanded to give rise to the knockdown line. Successful knockdown was validated by qRT-PCR (>80% E RN1 knockdown).

### Immunofluorescence and confocal microscopy

hHOs were transferred to microcentrifuge tubes (Eppendorf) using a wide bore 200 mL pipette tip to avoid disruption to the organoids and fixed in 4% paraformaldehyde solution. Fixation was followed by washes in phosphate-buffered saline (PBS)-Glycine (20 mM) and incubation in blocking/permeabilization solution containing 10% Donkey Normal Serum, 0.5% Triton X-100, 0.5% bovine serum albumin (BSA) in PBS on a thermal mixer (Thermo Scientific) at minimum speed at 4°C overnight. hHOs were then washed 3 times in PBS and incubated with primary antibodies in Antibody Solution (1% Donkey Normal Serum, 0.5% Triton X-100, 0.5% BSA in PBS) on a thermal mixer at minimum speed at 4°C for 24 hours. Primary antibody exposure was followed by 3 washes in PBS and incubation with secondary antibodies in Antibody Solution on a thermal mixer at minimum speed at 4°C for 24 hours in the dark. stained hHOs were washed 3 times in PBS before being mounted on glass microscope slides (Fisher Scientific) using Vectashield Vibrance Antifade Mounting Medium (Vector Laboratories). Samples were imaged using confocal laser scanning microscopy (Nikon Instruments A1 Confocal Laser Microscope). Images were analyzed using Fiji (https://imagej.net/Fiji).

### qRT-PCR and gene expression analysis

RNA was extracted from organoids at each condition/time point using the RNeasy Mini Kit (Qiagen). Once extracted, RNA was quantified using a NanoDrop (Mettler Toledo), with a concentration of at least 20 ng/µL being required to proceed with reverse transcription. cDNA was synthesized using the Quantitect Reverse Transcription Kit (Qiagen) and stored at -20°C for further use. Primers for qRT-PCR were designed using the Primer Quest tool (Integrated DNA Technologies) and SYBR Green (Qiagen) was used as the DNA intercalating dye. qRT-PCR plates were run using the QuantStudio 5 Real-Time PCR system (Applied Biosystems) with a total reaction volume of 20 µL. Expression levels of genes of interest were normalized to *HPRT1* levels and fold change values were obtained using the 2-ΔΔCT method. At least 3 independent samples were run for each gene expression assay.

### Organoid dissociation

Organoids were dissociated into a single-celled suspension using a modified protocol using the STEMdiff Cardiomyocyte Dissociation Kit (STEMCELL Technologies). Upon being transferred to a microcentrifuge tube, organoids were washed with PBS, submerged in 200 μL of warm dissociation media (37 °C), and placed on a thermal mixer at 37 °C and 300rpm for 5 minutes. Then, the supernatant was collected and transferred to a 15 mL falcon tube (Corning) containing 5mL of respective media (Control, MM, EMM1, etc.) containing 2% BSA (Thermo Fisher Scientific). An additional 200 μL of warm dissociation media (37 °C) was then added back to the organoid on a thermal mixer (37 °C). The organoid dissociation media solution was then pipetted up and down gently 3-5 times. The organoid was allowed to sit on the thermal mixer for an additional 5 minutes. If the organoid remained visible, the process was repeated. Once the organoid was no longer visible, the microcentrifuge tube solution was pipetted up and down gently 3-5 times and its entire contents were transferred to the 15 mL falcon tube containing the respective media + 2% BSA and cells. These tubes were then centrifuged at 300 g for 5 minutes. The supernatant was aspirated, and the cell pellets were resuspended in respective media + 2% BSA. A hemocytometer was used to determine viability and cell counts.

### Cardiomyocyte hypertrophy quantification

Hypertrophy was determined in single cell cardiomyocyte cultures produced from organoid dissociation 24 hours after plating. Organoids were stained with anti-cTnT antibodies for cytoplasm visualization and counterstained with DAPI as described under immunofluorescence methods. Cell hypertrophy was quantified using histomorphometry tools in ImageJ/Fiji.

### Mitochondrial staining

mitochondrial status within human heart organoids was visualized using Mitotracker Deep Red FM (Thermo Fisher Scientific). Mitotracker was prepared according to the manufacturer’s instructions. Organoids were washed twice using RPMI 1640 basal medium, then Mitotracker was added at a final concentration of 100 nM and incubated for 30 minutes at 37 °C and 5% CO2. Organoids were then washed twice and transferred to a chambered coverglass slide (Cellvis) using a cut 200 μL pipette tip. Images were acquired using a Cellvivo microscope (Olympus). Data was processed using ImageJ/Fiji.

### Single cell RNA-sequencing

Sequencing of the 10x Genomics Single Cell 3’ Gene Expression library was prepared from the cells dissociated from four pooled organoids per condition at day 15 of hHO differentiation. The libraries were prepared using the 10x Chromium Next GEM Single Cell 3’ Kit, v3.1 and associated components. Completed libraries were QC’d and quantified using a combination of Qubit dsDNA HS, Agilent 4200 TapeStation HS DNA1000 and Invitrogen Collibri Library Quantification qPCR assays. The library was loaded onto an Illumina NextSeq 500 v1.5 Mid Output flow cell. Sequencing was performed in a custom paired end format: 28 bases for read 1 which captures the 10x cell barcode and Unique Molecular Identifier (UMI), and 90 bases for read 2 which is the RNA portion of the library fragment. Base calling was done by Illumina Real Time Analysis (RTA) v2.4.11 and output of RTA was demultiplexed and converted to FastQ format with Illumina Bcl2fastq v2.20.0. After demultiplexing and FastQ conversion, secondary analysis was performed using cellranger count (v6.0.0). Analysis was executed using 10X Loupe Browser 6 for k-means clustering and t-SNE visualization, Enrichr (https://maayanlab.cloud/Enrichr/) for gene ontologies and DiVenn 2.0 (https://divenn.tch.harvard.edu/) for differentially expressed genes analysis.

### Calcium imaging

Calcium transients were observed in heart organoids by live imaging of organoids stained with Fluo-4 (Invitrogen) sing a super resolution microscope (Olympus CellVivo). Fluo-4 staining was conducted according to manufacturer’s instructions. Briefly, Fluo- 4-AM was reconstituted in DMSO to a final stock solution concentration of 0.5 mM. Fluo-4-AM. 1 µM of Fluo-4-AM was added directly to the organoid well and incubated for 30 minutes. The organoid was then washed twice in culture media and taken to the imaging microscope immediately. Organoids were transferred to a chambered cover glass in 100 µL of media. Several 10 seconds recordings were acquired for each organoid at 50 frames per second. Fluorescence intensity change was expressed as Δ*F*/*F*_0_ and was obtained analyzing videos using ImageJ/Fiji.

### ROS imaging

Organoids were stained with CellROX Green according to manufacturer’s instructions. Briefly, CellROX Green was added to the well of individual organoids at a final concentration of 5 µM and incubated for 30 minutes at 37°C, 5% CO_2_. Organoids were then washed twice in fresh culture medium and imaged immediately in an Olympus cellVivo microscope.

### LC-MS lipidomics

For organoid samples, at day 15, 10 organoids per condition were placed in individual tubes without any media, and flash frozen in a -80°C freezer. For medium, supernatants were placed in individual Eppendorf tubes and flash frozen in a -80°C freeze after a short 2-min 300 g centrifugation step to remove debris and other cellular remnants. Sample metabolite extraction and solid phase extraction (SPE) sample cleanup followed methods described before^79^. Briefly 100 µL of media (thawed on ice) or individual organoid samples (on dry ice) were spiked with 5 ng of d8-arachidonic acid. To each organoid sample, 250 µL of - 20°C chilled 75% ethanol was added, and samples were homogenized in a bead mill for 2 minutes. Homogenates were transferred to 2.0 mL centrifuge tubes, and an additional 750 µL of -20°C chilled 75% ethanol was added. Samples were vortexed for 30 minutes, then incubated at -20°C for one hour to precipitate proteins. Samples were then centrifuged at 15,000 x g for 20 minutes. The supernatants were transferred to new 2.0 mL centrifuge tubes, and the remaining protein pellets were re-extracted and the supernatants pooled with those from the first extraction. Media samples were processed similarly using a 3:1 ratio of ethanol:media, with omission of the homogenization step. The pooled supernatants were diluted with 1.0 mL of HPLC water and applied to preconditioned SPE columns (Phenomenex Strata-X polymeric reverse phase, 10 mg/1 mL, 33 µm) using vacuum assisted pull-through. The flow-through was discarded, and the SPE columns were washed with 900 µL of 10% methanol, then eluted with 400 µL of 100% ethanol. Samples were dried under vacuum in a speedvac centrifuge, reconstituted in 50 µL of acetonitrile by vortexing for 15 minutes, then transferred to LC-MS vials containing small volume inserts. Samples were stored at -80°C until analysis. The LC-MS platform consisted of a Shimadzu Prominence HPLC coupled to a Thermo LTQ-Orbitrap Velos mass spectrometer. The HPLC column was a Phenomenex 2.0 mmx150 mm Synergi HydroRP- C18 (4 µm, 80 Angstrom pore size) equipped with a guard cartridge of the same column chemistry. The LC gradient consisted of solvent A, 70:30 water:acetonitrile (v:v) containing 0.1% acetic acid, solvent B, which was 50:50 isopropanol:acetonitrile containing 0.02% acetic acid. The flow rate was 200 µL per minute and the column oven was held at 45 C. The autosampler was held at 4 C. 10 µL of each sample was injected. The gradient conditions used were: Time 0-2 minutes, 1% solvent B. Column eluant was diverted to waste using a 2-position 6 port valve. At time=2.0 minutes, Solvent B was increased to 50%, and a linear gradient from 50% to 65% B was run between 2.0 and 10 minutes. Solvent B then increased linearly to 99% B between 10 and 16 minutes, and solvent B was held constant at 99% until 24 minutes. Solvent B was then returned to 1 to re-equilibrate the column for 5 minutes. Column eluent was introduced to a Thermo LTQ-Orbitrap Velos mass spectrometer via a heated electrospray ionization source. The mass spectrometer was operated in negative ion mode at 30,000 resolution with full scan MS data collected from 200-700 m/z. Data-dependent product ion spectra were collected on the 4 most abundant ions at 7,500 resolution using the FT analyzer. The electrospray ionization source was maintained at a spray voltage of 4.5 kV with sheath gas at 30 (arbitrary units), auxiliary gas at 10 (arbitrary units) and sweep gas at 2.0 (arbitrary units). The inlet of the mass spectrometer was held at 350°C, and the S-lens was set to 35%. The heated ESI source was maintained at 350°C. For data analysis, Chromatographic alignment, isotope correction, peak identification and peak area calculations were performed using MAVEN software. Concentrations of each analyte were determined against the peak area of the internal standard (D8-arachidonic acid). Confirmed analytes were identified by comparison against authentic reference standards.

### ELISA assays

Heart organoids total proteins were extracted from 8 individual organoids per condition using M-PER mammalian protein extract reagent (Thermo). Total proteins concentration in lysates was measured by BCA Protein Assay Kit (Thermo). FADS2 protein in heart organoids lysates was measured using a human FADS2 RTU ELISA Kit (MyBioSource).

### RNA structure analysis

Geneious Prime was used for analysis of mRNA sequences and secondary structure visualization of all RNA candidates.

### Statistics and Reproducibility

All analyses were performed using Excel and GraphPad software. Statistical significance was evaluated with a standard unpaired Student t-test or one-way ANOVA as appropriate. All data are presented as mean ± S.D. and represent a minimum of 3 independent experiments with at least 3 technical replicates per experiment unless otherwise stated.

### Data Availability

The organoid RNA-Sequencing data sets have been deposited in the National Center for Biotechnology Information Gene Expression Omnibus repository under accession code GSE201343. All other data generated and/or analyzed in this study are provided in the published article and its supplementary information files or from the corresponding author upon request.

## Acknowledgments

We wish to thank the MSU Mass Spectrometry Core, the MSU Genomics Core, the MSU Advanced Imaging Core and the IQ Microscopy Core. Work in Dr. Aguirre’s laboratory was supported by the NIH under award numbers K01HL135464, R01HL151505, by a Pilot and Feasibility Grant from the Michigan Diabetes Research Center (NIH Grant P30-DK020572), by the American Heart Association under award number 19IPLOI34660342, and by the Spectrum-MSU Alliance Foundation. We also wish to thank all members of the Aguirre Lab for their valuable comments and advice.

## Author Contributions

AK, YLI, MA and AA designed experiments and conceptualized the work. AK and YLI performed all experiments and data analysis. MA, MAG, HL and AW performed cell and organoid culture and qPCR. BV performed live calcium recordings. TL performed LC-MS and data analysis. ARH performed qPCR and lentivirus experiments. AK, YLI, IO, CC and AA wrote the manuscript and provided valuable suggestions and advice. AA supervised all the work.

## REFERENCES

1. Zaidi, S., and Brueckner, M. (2017). Genetics and Genomics of Congenital Heart Disease. Circulation research 120, 923–940. 10.1161/CIRCRESAHA.116.309140.

2. Oyen, N., Diaz, L.J., Leirgul, E., Boyd, H.A., Priest, J., Mathiesen, E.R., Quertermous, T., Wohlfahrt, J., and Melbye, M. (2016). Prepregnancy Diabetes and Offspring Risk of Congenital Heart Disease: A Nationwide Cohort Study. Circulation 133, 2243–2253. 10.1161/CIRCULATIONAHA.115.017465.

3. Basu, M., and Garg, V. (2018). Maternal hyperglycemia and fetal cardiac development: Clinical impact and underlying mechanisms. Birth Defects Res 110, 1504–1516. 10.1002/bdr2.1435.

4. Ornoy, A., Reece, E.A., Pavlinkova, G., Kappen, C., and Miller, R.K. (2015). Effect of maternal diabetes on the embryo, fetus, and children: congenital anomalies, genetic and epigenetic changes and developmental outcomes. Birth Defects Res C Embryo Today 105, 53–72. 10.1002/bdrc.21090.

5. Tabib, A., Shirzad, N., Sheikhbahaei, S., Mohammadi, S., Qorbani, M., Haghpanah, V., Abbasi, F., Hasani-Ranjbar, S., and Baghaei-Tehrani, R. (2013). Cardiac malformations in fetuses of gestational and pre gestational diabetic mothers. Iran J Pediatr 23, 664–668.

6. Cho, N.H., Shaw, J.E., Karuranga, S., Huang, Y., da Rocha Fernandes, J.D., Ohlrogge, A.W., and Malanda, B. (2018). IDF Diabetes Atlas: Global estimates of diabetes prevalence for 2017 and projections for 2045. Diabetes Res Clin Pract 138, 271–281. 10.1016/j.diabres.2018.02.023.

7. Wu, Y., Liu, B., Sun, Y., Du, Y., Santillan, M.K., Santillan, D.A., Snetselaar, L.G., and Bao, W. (2020). Association of Maternal Prepregnancy Diabetes and Gestational Diabetes Mellitus With Congenital Anomalies of the Newborn. Diabetes Care 43, 2983–2990. 10.2337/dc20-0261.

8. Peng, T.Y., Ehrlich, S.F., Crites, Y., Kitzmiller, J.L., Kuzniewicz, M.W., Hedderson, M.M., and Ferrara, A. (2017). Trends and racial and ethnic disparities in the prevalence of pregestational type 1 and type 2 diabetes in Northern California: 1996-2014. Am J Obstet Gynecol 216, 177 e171–177 e178. 10.1016/j.ajog.2016.10.007.

9. Buchanan, M., Thomas A, and Kitzmiller, M., John L (1994). Metabolic interactions of diabetes and pregnancy. Annual review of medicine 45, 245–260.

10. Correa, A. (2016). Pregestational diabetes mellitus and congenital heart defects. Am Heart Assoc.

11. Engineer, A., Saiyin, T., Lu, X., Kucey, A.S., Urquhart, B.L., Drysdale, T.A., Norozi, K., and Feng, Q. (2018). Sapropterin Treatment Prevents Congenital Heart Defects Induced by Pregestational Diabetes Mellitus in Mice. Journal of the American Heart Association 7, e009624. 10.1161/JAHA.118.009624.

12. Lehtoranta, L., Koskinen, A., Vuolteenaho, O., Laine, J., Kytö, V., Soukka, H., Ekholm, E., and Räsänen, J. (2017). Gestational hyperglycemia reprograms cardiac gene expression in rat offspring. Pediatric Research 82, 356–361.

13. Lehtoranta, L., Vuolteenaho, O., Laine, V.J., Koskinen, A., Soukka, H., Kytö, V., Määttä, J., Haapsamo, M., Ekholm, E., and Räsänen, J. (2013). Maternal hyperglycemia leads to fetal cardiac hyperplasia and dysfunction in a rat model. American Journal of Physiology-Endocrinology and Metabolism 305, E611–E619.

14. Piddington, R., Joyce, J., Dhanasekaran, P., and Baker, L. (1996). Diabetes mellitus affects prostaglandin E 2 levels in mouse embryos during neurulation. Diabetologia 39, 915–920.

15. Wang, F., Reece, E.A., and Yang, P. (2015). Oxidative stress is responsible for maternal diabetes-impaired transforming growth factor beta signaling in the developing mouse heart. Am J Obstet Gynecol 212, 650 e651–611. 10.1016/j.ajog.2015.01.014.

16. Wu, Y., Reece, E.A., Zhong, J., Dong, D., Shen, W.-B., Harman, C.R., and Yang, P. (2016). Type 2 diabetes mellitus induces congenital heart defects in murine embryos by increasing oxidative stress, endoplasmic reticulum stress, and apoptosis. American journal of obstetrics and gynecology 215, 366. e361–366. e310.

17. Gupta, A., Chacko, V.P., Schär, M., Akki, A., and Weiss, R.G. (2011). Impaired ATP kinetics in failing in vivo mouse heart. Circulation: Cardiovascular Imaging 4, 42–50.

18. Hamlin, R.L., and Altschuld, R.A. (2011). Extrapolation from mouse to man. Am Heart Assoc.

19. Loiselle, D.S., and Gibbs, C.L. (1979). Species differences in cardiac energetics. Am J Physiol 237, H90–98. 10.1152/ajpheart.1979.237.1.H90.

20. Kretzschmar, K., and Clevers, H. (2016). Organoids: modeling development and the stem cell niche in a dish. Developmental cell 38, 590–600.

21. Paşca, S.P. (2019). Assembling human brain organoids. Science 363, 126–127.

22. Rossi, G., Broguiere, N., Miyamoto, M., Boni, A., Guiet, R., Girgin, M., Kelly, R.G., Kwon, C., and Lutolf, M.P. (2021). Capturing cardiogenesis in gastruloids. Cell stem cell 28, 230–240. e236.

23. Sasai, Y. (2013). Next-generation regenerative medicine: organogenesis from stem cells in 3D culture. Cell stem cell 12, 520–530.

24. Weeber, F., Ooft, S.N., Dijkstra, K.K., and Voest, E.E. (2017). Tumor organoids as a pre- clinical cancer model for drug discovery. Cell chemical biology 24, 1092–1100.

25. Bredenoord, A.L., Clevers, H., and Knoblich, J.A. (2017). Human tissues in a dish: the research and ethical implications of organoid technology. Science 355, eaaf9414.

26. Lewis-Israeli, Y.R., Wasserman, A.H., and Aguirre, A. (2021). Heart organoids and engineered heart tissues: Novel tools for modeling human cardiac biology and disease. Biomolecules 11, 1277.

27. Tuveson, D., and Clevers, H. (2019). Cancer modeling meets human organoid technology. Science 364, 952–955.

28. Hudson, J., Mills, R., Titmarsh, D., Koenig, X., Parker, B., Ryall, J., Quaife-Ryan, G., Voges, H., Hodson, M., and Ferguson, C. (2017). Functional screening in human cardiac organoids reveals a metabolic mechanism for cardiomyocyte cell cycle arrest. Heart, Lung and Circulation 26, S207–S208.

29. Lewis-Israeli, Y.R., Wasserman, A.H., Gabalski, M.A., Volmert, B.D., Ming, Y., Ball, K.A., Yang, W., Zou, J., Ni, G., and Pajares, N. (2021). Self-assembling human heart organoids for the modeling of cardiac development and congenital heart disease. Nature communications 12, 5142.

30. Volmert, B., Riggs, A., Wang, F., Juhong, A., Kiselev, A., Kostina, A., O’hern, C., Muniyandi, P., Wasserman, A., and Huang, A. (2022). A patterned human heart tube organoid model generated by pluripotent stem cell self-assembly. bioRxiv, 2022.2012. 2016.519611.

31. Hofbauer, P., Jahnel, S.M., Papai, N., Giesshammer, M., Deyett, A., Schmidt, C., Penc, M., Tavernini, K., Grdseloff, N., and Meledeth, C. (2021). Cardioids reveal self-organizing principles of human cardiogenesis. Cell 184, 3299–3317. e3222.

32. Guemes, M., Rahman, S.A., and Hussain, K. (2016). What is a normal blood glucose? Arch Dis Child 101, 569–574. 10.1136/archdischild-2015-308336.

33. Association, A.D. (2009). Diagnosis and classification of diabetes mellitus. Diabetes care 32, S62.

34. Al-Biltagi, M., and El Amrousy, D. (2021). Cardiac changes in infants of diabetic mothers. World Journal of Diabetes 12, 1233.

35. Depla, A., De Wit, L., Steenhuis, T., Slieker, M., Voormolen, D., Scheffer, P., De Heus, R., Van Rijn, B., and Bekker, M. (2021). Effect of maternal diabetes on fetal heart function on echocardiography: systematic review and meta-analysis. Ultrasound in Obstetrics & Gynecology 57, 539–550.

36. Hornberger, L.K. (2006). Maternal diabetes and the fetal heart. BMJ Publishing Group Ltd.

37. Reinking, B.E., Wedemeyer, E.W., Weiss, R.M., Segar, J.L., and Scholz, T.D. (2009). Cardiomyopathy in offspring of diabetic rats is associated with activation of the MAPK and apoptotic pathways. Cardiovascular diabetology 8, 1–9.

38. Schwartz, R., Gruppuso, P.A., Petzold, K., Brambilla, D., Hiilesmaa, V., and Teramo, K.A. (1994). Hyperinsulinemia and macrosomia in the fetus of the diabetic mother. Diabetes care 17, 640–648.

39. Giacco, F., and Brownlee, M. (2010). Oxidative stress and diabetic complications. Circulation research 107, 1058–1070.

40. Wang, S., Binder, P., Fang, Q., Wang, Z., Xiao, W., Liu, W., and Wang, X. (2018). Endoplasmic reticulum stress in the heart: insights into mechanisms and drug targets. British journal of pharmacology 175, 1293–1304.

41. Maurel, M., Chevet, E., Tavernier, J., and Gerlo, S. (2014). Getting RIDD of RNA: IRE1 in cell fate regulation. Trends Biochem Sci 39, 245–254. 10.1016/j.tibs.2014.02.008.

42. Valentine, R., and Valentine, D. Progr. Lipid Res.–2004. V 43, 383–402.

43. Asatryan, A., and Bazan, N.G. (2017). Molecular mechanisms of signaling via the docosanoid neuroprotectin D1 for cellular homeostasis and neuroprotection. Journal of Biological Chemistry 292, 12390–12397.

44. Li, G., Li, Y., Xiao, B., Cui, D., Lin, Y., Zeng, J., Li, J., Cao, M.-J., and Liu, J. (2021). Antioxidant activity of docosahexaenoic acid (DHA) and its regulatory roles in mitochondria. Journal of Agricultural and Food Chemistry 69, 1647–1655.

45. Zhang, Q., Schepis, A., Huang, H., Yang, J., Ma, W., Torra, J., Zhang, S.-Q., Yang, L., Wu, H., and Nonell, S. (2019). Designing a green fluorogenic protease reporter by flipping a beta strand of GFP for imaging apoptosis in animals. Journal of the American Chemical Society 141, 4526–4530.

46. Hollien, J., Lin, J.H., Li, H., Stevens, N., Walter, P., and Weissman, J.S. (2009). Regulated Ire1-dependent decay of messenger RNAs in mammalian cells. J Cell Biol 186, 323–331. 10.1083/jcb.200903014.

47. Vaittinen, M., Walle, P., Kuosmanen, E., Männistö, V., Käkelä, P., Ågren, J., Schwab, U., and Pihlajamäki, J. (2016). FADS2 genotype regulates delta-6 desaturase activity and inflammation in human adipose tissue. Journal of lipid research 57, 56–65.

48. Brown, K.M., Sharma, S., Baker, E., Hawkins, W., van der Merwe, M., and Puppa, M.J. (2019). Delta-6-desaturase (FADS2) inhibition and omega-3 fatty acids in skeletal muscle protein turnover. Biochem Biophys Rep 18, 100622. 10.1016/j.bbrep.2019.100622.

49. Le, C.H., Mulligan, C.M., Routh, M.A., Bouma, G.J., Frye, M.A., Jeckel, K.M., Sparagna, G.C., Lynch, J.M., Moore, R.L., and McCune, S.A. (2014). Delta-6-desaturase links polyunsaturated fatty acid metabolism with phospholipid remodeling and disease progression in heart failure. Circulation: Heart Failure 7, 172–183.

50. Arshad, Z., Rezapour-Firouzi, S., Ebrahimifar, M., Jarrahi, A.M., and Mohammadian, M. (2019). Association of delta-6-desaturase expression with aggressiveness of cancer, diabetes mellitus, and multiple sclerosis: a narrative review. Asian Pacific Journal of Cancer Prevention: APJCP 20, 1005.

51. Lewis-Israeli, Y.R., Abdelhamid, M., Olomu, I., and Aguirre, A. (2022). Modeling the Effects of Maternal Diabetes on the Developing Human Heart Using Pluripotent Stem Cell-Derived Heart Organoids. Curr Protoc 2, e461. 10.1002/cpz1.461.

52. Narchi, H., and Kulaylat, N. (2000). Heart disease in infants of diabetic mothers. Images Paediatr Cardiol 2, 17–23.

53. Kc, K., Shakya, S., and Zhang, H. (2015). Gestational diabetes mellitus and macrosomia: a literature review. Annals of Nutrition and Metabolism 66, 14–20.

54. Wende, A.R., and Abel, E.D. (2010). Lipotoxicity in the heart. Biochimica et Biophysica Acta (BBA)-Molecular and Cell Biology of Lipids 1801, 311–319.

55. Engineer, A., Saiyin, T., Greco, E.R., and Feng, Q. (2019). Say NO to ROS: Their roles in embryonic heart development and pathogenesis of congenital heart defects in maternal diabetes. Antioxidants 8, 436.

56. Moazzen, H., Lu, X., Ma, N.L., Velenosi, T.J., Urquhart, B.L., Wisse, L.J., Gittenberger-de Groot, A.C., and Feng, Q. (2014). N-Acetylcysteine prevents congenital heart defects induced by pregestational diabetes. Cardiovascular diabetology 13, 1–13.

57. Kumar, S.D., Vijaya, M., Samy, R.P., Dheen, S.T., Ren, M., Watt, F., Kang, Y.J., Bay, B.-H., and Tay, S.S.W. (2012). Zinc supplementation prevents cardiomyocyte apoptosis and congenital heart defects in embryos of diabetic mice. Free Radical Biology and Medicine 53, 1595–1606.

58. Kumar, S.D., Dheen, S.T., and Tay, S.S.W. (2007). Maternal diabetes induces congenital heart defects in mice by altering the expression of genes involved in cardiovascular development. Cardiovascular diabetology 6, 1–14.

59. Gandhi, J.A., Zhang, X.Y., and Maidman, J.E. (1995). Fetal cardiac hypertrophy and cardiac function in diabetic pregnancies. American journal of obstetrics and gynecology 173, 1132–1136.

60. Zhong, J., Xu, C., Gabbay-Benziv, R., Lin, X., and Yang, P. (2016). Superoxide dismutase 2 overexpression alleviates maternal diabetes-induced neural tube defects, restores mitochondrial function and suppresses cellular stress in diabetic embryopathy. Free Radical Biology and Medicine 96, 234–244.

61. Alam, M.J., Uppulapu, S.K., Tiwari, V., Varghese, B., Mohammed, S.A., Adela, R., Arava, S.K., and Banerjee, S.K. (2022). Pregestational diabetes alters cardiac structure and function of neonatal rats through developmental plasticity. Frontiers in Cardiovascular Medicine.

62. Jia, G. (2018). Hill, and JR Sowers, Diabetic Cardiomyopathy: An Update of Mechanisms Contributing to This Clinical Entity. Circulation research 122, 624–638.

63. Quijada, P., Trembley, M.A., and Small, E.M. (2020). The Role of the Epicardium During Heart Development and Repair. Circulation research 126, 377–394. 10.1161/CIRCRESAHA.119.315857.

64. Geyik, F., Ozuynuk, A., Erkan, A., Ekici, B., and Coban, N. (2020). Association of itln1 gene polymorphism in coronary artery disease with type 2 diabetes. Atherosclerosis 315, e127.

65. Au-Yeung, C.-L., Yeung, T.-L., Achreja, A., Zhao, H., Yip, K.-P., Kwan, S.-Y., Onstad, M., Sheng, J., Zhu, Y., and Baluya, D.L. (2020). ITLN1 modulates invasive potential and metabolic reprogramming of ovarian cancer cells in omental microenvironment. Nature communications 11, 3546.

66. Cui, X., Zhang, Y., Lu, Y., and Xiang, M. (2022). ROS and endoplasmic reticulum stress in pulmonary disease. Frontiers in Pharmacology 13.

67. Moncan, M., Mnich, K., Blomme, A., Almanza, A., Samali, A., and Gorman, A.M. (2021). Regulation of lipid metabolism by the unfolded protein response. Journal of cellular and molecular medicine 25, 1359–1370.

68. Ho, N., Xu, C., and Thibault, G. (2018). From the unfolded protein response to metabolic diseases–lipids under the spotlight. Journal of cell science 131, jcs199307.

69. Travers, K.J., Patil, C.K., Wodicka, L., Lockhart, D.J., Weissman, J.S., and Walter, P. (2000). Functional and genomic analyses reveal an essential coordination between the unfolded protein response and ER-associated degradation. Cell 101, 249–258.

70. Halbleib, K., Pesek, K., Covino, R., Hofbauer, H.F., Wunnicke, D., Hänelt, I., Hummer, G., and Ernst, R. (2017). Activation of the unfolded protein response by lipid bilayer stress. Molecular cell 67, 673–684. e678.

71. Volmer, R., van der Ploeg, K., and Ron, D. (2013). Membrane lipid saturation activates endoplasmic reticulum unfolded protein response transducers through their transmembrane domains. Proceedings of the National Academy of Sciences 110, 4628–4633.

72. Nath, A., Oak, A., Chen, K.Y., Li, I., Splichal, R.C., Portis, J., Foster, S., Walton, S.P., and Chan, C. (2021). Palmitate-Induced IRE1–XBP1–ZEB Signaling Represses Desmoplakin Expression and Promotes Cancer Cell MigrationPalmitate Induced DSP Loss via IRE1–XBP1–ZEB. Molecular Cancer Research 19, 240–248.

73. Manivannan, S., Mansfield, C., Zhang, X., Kodigepalli, K.M., Majumdar, U., Garg, V., and Basu, M. (2021). Single-cell transcriptomic profiling unveils cardiac cell-type specific response to maternal hyperglycemia underlying the risk of congenital heart defects. BioRxiv, 2021.2005. 2028.446177.

74. Richards, D.J., Li, Y., Kerr, C.M., Yao, J., Beeson, G.C., Coyle, R.C., Chen, X., Jia, J., Damon, B., and Wilson, R. (2020). Human cardiac organoids for the modelling of myocardial infarction and drug cardiotoxicity. Nature Biomedical Engineering 4, 446–462.

75. Filippo Buono, M., von Boehmer, L., Strang, J., P. Hoerstrup, S., Y. Emmert, M., and Nugraha, B. (2020). Human cardiac organoids for modeling genetic cardiomyopathy. Cells 9, 1733.

76. Binder, P., Wang, S., Radu, M., Zin, M., Collins, L., Khan, S., Li, Y., Sekeres, K., Humphreys, N., and Swanton, E. (2019). Pak2 as a novel therapeutic target for cardioprotective endoplasmic reticulum stress response. Circulation research 124, 696–711.

77. Lee, R., Xu, B., Rame, J.E., Felkin, L.E., Barton, P., and Dries, D.L. (2014). Regulated inositol-requiring protein 1-dependent decay as a mechanism of corin RNA and protein deficiency in advanced human systolic heart failure. Journal of the American Heart Association 3, e001104.

78. Basak, S., Mallick, R., and Duttaroy, A.K. (2020). Maternal docosahexaenoic acid status during pregnancy and its impact on infant neurodevelopment. Nutrients 12, 3615.

79. Watrous, J.D., Niiranen, T.J., Lagerborg, K.A., Henglin, M., Xu, Y.-J., Rong, J., Sharma, S., Vasan, R.S., Larson, M.G., and Armando, A. (2019). Directed non-targeted mass spectrometry and chemical networking for discovery of eicosanoids and related oxylipins. Cell chemical biology 26, 433–442. e434.

