## Supplementary data for "ER stress and lipid imbalance drive embryonic cardiomyopathy in a human heart organoid model of pregestational diabetes"

**SUPPLEMENTARY FIGURES**

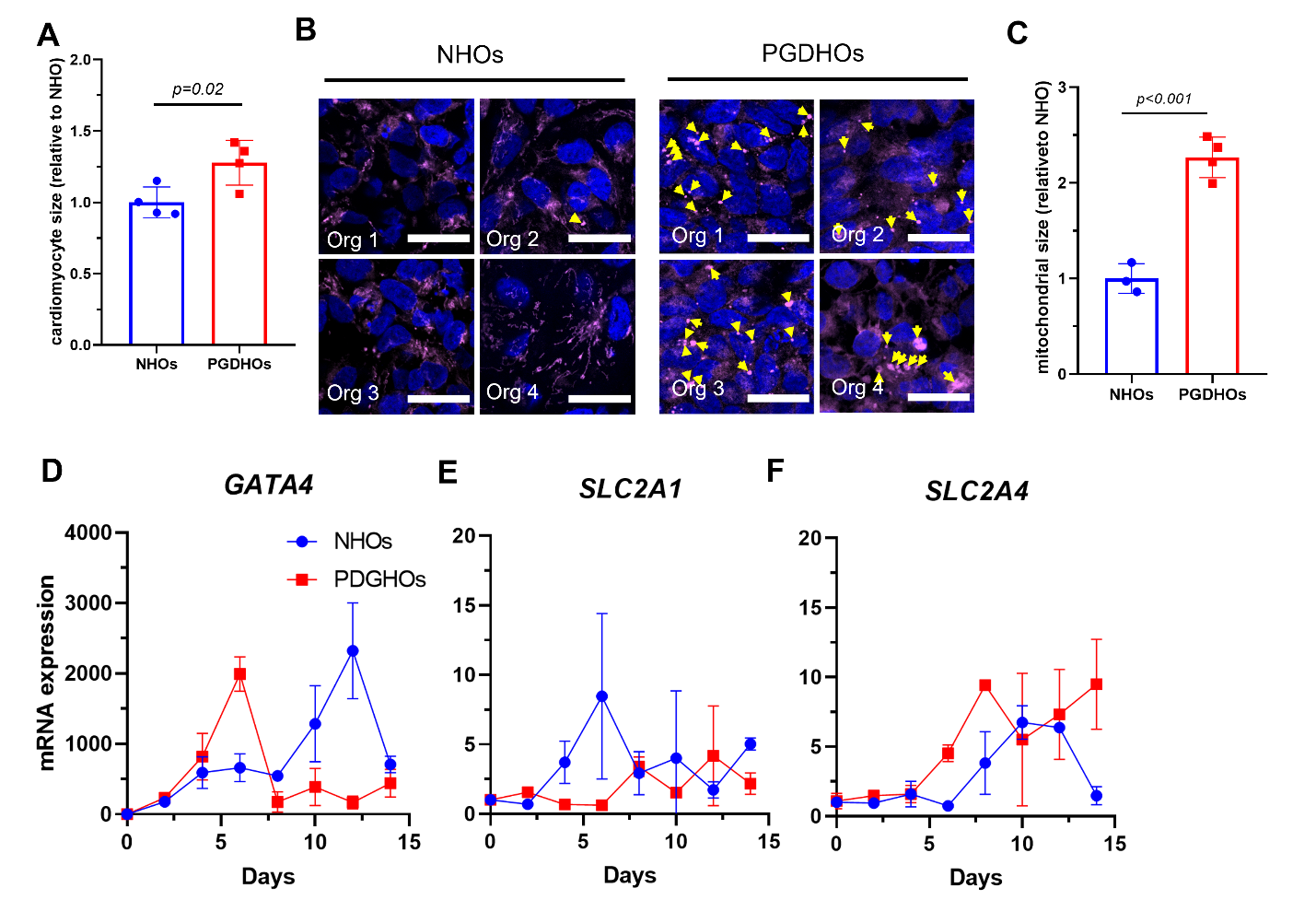

**Supp. Fig. 1**. **A**, Quantification of cardiomyocyte area compared to NHOs; n=4 organoids, n=145 cardiomyocytes for NHOs, n=193 cardiomyocytes for PGDHOs; value = mean ± SD, unpaired t-tests. **B**, Immunofluorescence image analysis showing mitochondrial staining from 4 independent organoids, arrowheads indicated mitochondrial swelling; scale bar = 25 µm. **C**, Quantification of mitochondrial swelling compared to NHOs; n=4; value = mean ± SD, unpaired t-tests. **D**, Gene expression analysis time course comparison between NHOs and PGDHOs for GATA4 (**E**) the glucose transporter SLC2A1 and (**F**) the glucose transporter SLC2A4.

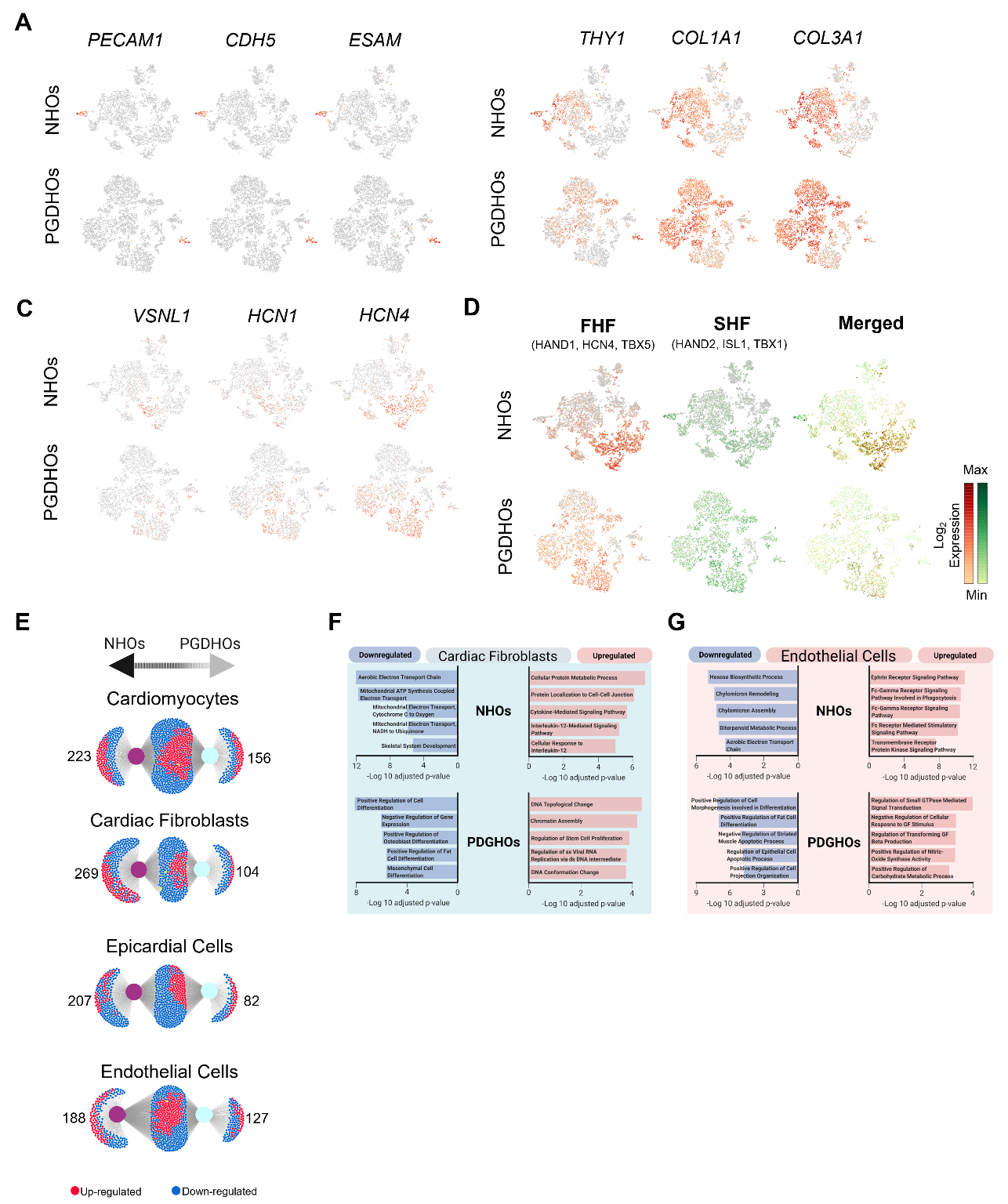

**Supp. Fig. 2**. **A**, t-SNE plots showing key markers for endothelial cells, (**B**) cardiac fibroblasts and (**C**) sinoatrial node cells from. **D**, t-SNE plots showing overlap of clusters expressing key markers of early heart fields. **E**, Venn diagrams of significantly expressed DEGs from cardiomyocyte, cardiac fibroblasts, epicardial cells, and endothelial cells clusters between day 15 NHOs and PGDHOs. **F**, **G** gene ontology analysis of biological processes associated with DEGs in the cardiac fibroblasts cluster (**F**) and the endothelial cells cluster (**G**).

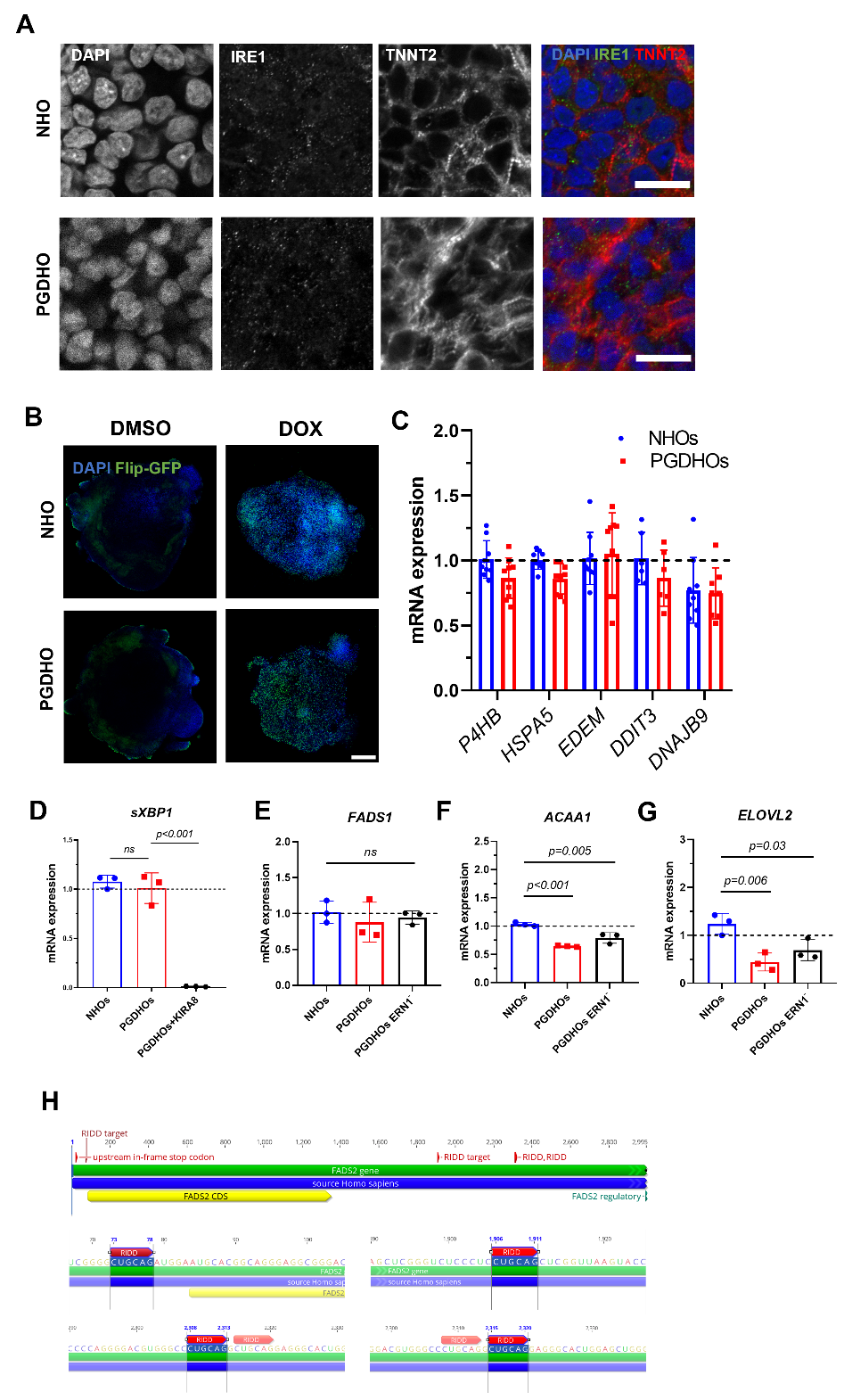

**Supp. Fig. 3**. **A,** Immunofluorescence images of day 14 NHOs and PGDHOs stained for unphosphorylated IRE1 (green), cardiomyocyte marker TNNT2 (red) and nuclear marker DAPI; n=7 organoids; scale bar=20 µm. **B,** Immunofluorescence images of organoids derived from a Flip-GFP transgenic iPSC line under control (NHO, top) and diabetic (PGDHO, bottom) conditions treated with vehicle (DMSO) or doxorubicin (positive control); scale bar = 200 µm. **C,** Gene expression analysis of UPR markers in NHOs and PGDHOs as determined by qRT-PCR; n=9 biological replicates of 3 pooled organoids; value = mean ± SD. **F**, qRT-PCR gene expression analysis of spliced *XBP1* in NHOs, PGDHOs; n=3 biological replicates of 3 pooled organoids; value = mean ± SD. **E-G**, Gene expression analysis of enzymes involved in VLCFA biosynthesis in NHOs, PGDHOs; n=3 biological replicates of 3 pooled organoids; value = mean ± SD. **H**, mRNA sequence of *FADS2* mRNA with marked consensus sites as potential targets for IRE1-RIDD degradation.
